# Conserved luminal C-terminal domain dynamically controls interdomain communication in sarcolipin

**DOI:** 10.1101/2020.03.28.013425

**Authors:** Rodrigo Aguayo-Ortiz, Eli Fernández-de Gortari, L. Michel Espinoza-Fonseca

## Abstract

Sarcolipin (SLN) mediates Ca^2+^ transport and metabolism in muscle by regulating the activity of the Ca^2+^ pump SERCA. SLN has a conserved luminal C-terminal domain that contributes to the its functional divergence among homologous SERCA regulators, but the precise mechanistic role of this domain remains poorly understood. We used all-atom molecular dynamics (MD) simulations of SLN totaling 77.5 μs to show that the N- (NT) and C-terminal (CT) domains function in concert. Analysis of the MD simulations showed that serial deletions of SLN C-terminus does not affect the stability of the peptide nor induce dissociation of SLN from the membrane but promotes a gradual decrease in both tilt angle of the transmembrane helix and the local thickness of the lipid bilayer. Mutual information analysis showed that the NT and CT domains communicate with each other in SLN, and that interdomain communication is partially or completely abolished upon deletion of the conserved segment Tyr29-Tyr31 as well as by serial deletions beyond this domain. Phosphorylation of SLN at residue Thr5 also induces changes in the communication between the CT and NT domains, thus providing additional evidence for interdomain communication within SLN. We found that interdomain communication is independent of the force field used and lipid composition, thus demonstrating that communication between the NT and CT domains is an intrinsic functional feature of SLN. We propose the novel hypothesis that the conserved C-terminus is an essential element required for dynamic control of SLN regulatory function.

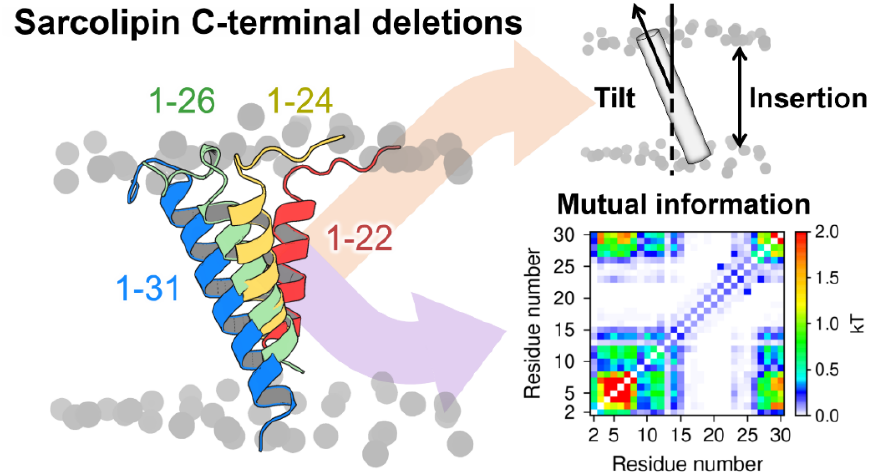

## 1. INTRODUCTION

Sarcolipin (SLN) is a 31-residue membrane protein found in the sarcoplasmic reticulum (SR) of cardiac atrial and skeletal muscle cells.^1^ SLN belongs to the family of proteins (which include phospholamban and myoregulin)^2,3^ that modulate the activity of the sarcoplasmic reticulum Ca^2+^-ATPase (SERCA), a ion-motive pump that clears cytosolic Ca^2+^ in muscle cells.^4^ SLN plays a key role in Ca^2+^ homeostasis of muscle cells by inhibiting SERCA, and also participates in metabolism by inducing unproductive ATPase activity of SERCA and stimulating non-shivering thermogenesis.^5^

SLN consists of three functional domains: (i) an unstructured N-terminus (NT, residues M1-E7) that contains a site for phosphorylation by the calmodulin-dependent protein kinase II (CaMKII) at residue T5;^6^ (ii) a transmembrane helix (TM, residues L8-V26) that is important for SERCA binding as well as for self-association;^7,8^ and (iii) a conserved luminal C-terminal domain (CT, residues R27-Y31) that protrudes into the SR lumen (**Figure 1**).^9,10^ Experiments and molecular simulation studies have shown that the NT domain plays a primary role in SERCA uncoupling, while the TM domain is necessary for binding and stabilizing the regulatory complex as well as for inhibiting Ca^2+^ affinity.^11–13^ However, the role of the highly conserved CT domain (**Figure S1**) is more elusive and difficult to interpret. Mutagenesis experiments using co-reconstituted proteoliposomes have suggested that the CT domain is essential for SERCA inhibition,^14^ and cell-based studies have shown that serial deletion of this domain impairs localization and retention in the endoplasmic reticulum (ER).^15^ While there is clear evidence for the functional roles of the NT and TM domains of SLN,^5^ there is less clear evidence on the functional role of the CT domain. In this study, we investigated the functional role of SLN’s CT domain using μs-long atomistic molecular dynamics (MD) simulations of wild-type SLN (SLN_WT_, residues M1-Y31) and nine serial deletions in the CT domain (SLN_1-22_ through SLN_1-30_) studied experimentally by Gramolini et al.^15^ Based on the results of these extensive simulations, we propose that the conserved C-terminus is an essential element required for dynamic control of SLN regulatory function in the cell.

**Figure 1.**
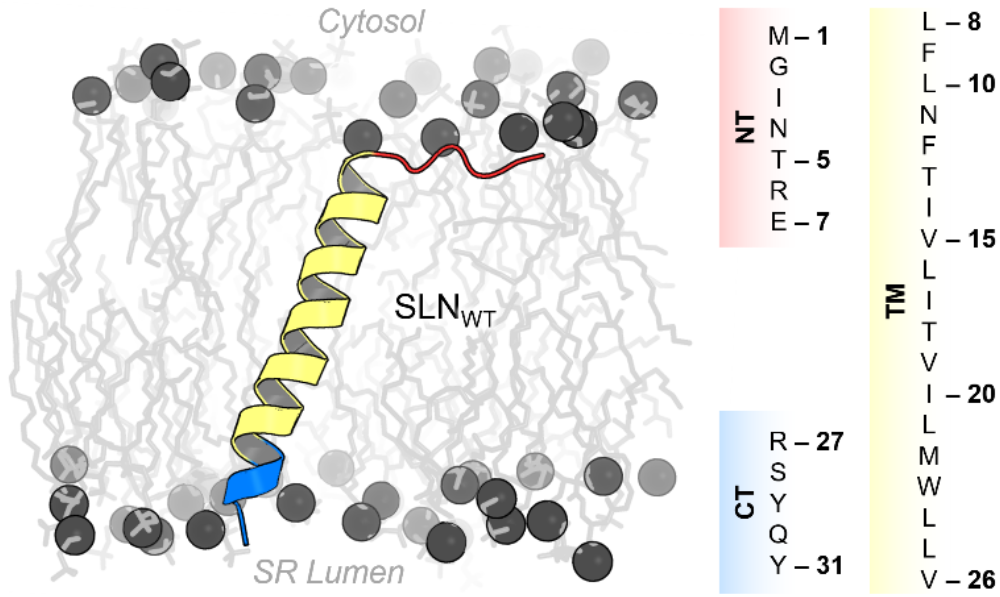
Structure and amino acid sequence of human SLN. The protein is shown as a ribbon representation and colored according to its functional regions: N-terminus (NT, red), transmembrane helix (TM, yellow), and luminal C-terminal (CT, blue) domains. The headgroup and tails of lipids are shown spheres and sticks, respectively. The columns on the right panel show the amino acid sequence for each functional region of SLN.

## 2. Methods

### 2.1. Setting up the SLN construct systems

Full-length NMR structures of sarcolipin (SLN) obtained from the Protein Data Bank (PDB: 1JDM^9^) were used as a starting point for the MD simulations. We performed a cluster structure analysis with a root mean square deviation (RMSD) threshold of 2.5 Å to select the six most representative SLN structures for simulation. Each structure obtained in this clustering analysis was used to generate ten SLN contructs of different amino acid lengths: 1-31 (SLN_WT_), 1-30 (SLN_1-30_), 1-29 (SLN_1-29_), 1-28 (SLN_1-28_), 1-27 (SLN_1-27_), 1-26 (SLN_1-26_), 1-25 (SLN_1-25_), 1-24 (SLN_1-24_), 1-23 (SLN_1-23_), and 1-22 (SLN_1-22_). An additional structural model of phosphoryated SLN was generated by modeling a phosphorylated threonine side chain at position 5. The spatial arrangement of SLN in a lipid membrane was calculated with the PPM web server.^16^ Each structure of the protein was inserted in a pre-equlibrated bilayer containing a total 160 molecules of 1-palmitoyl-2-oleoyl-sn-glycero-3-phosphocholine (POPC) lipids using the *membrane builder* module of CHARMM-GUI web server.^17,18^ The systems were solvated using a TIP3P water model with a minimum margin of 20 Å between the protein and the edges of the periodic box in the z-axis. Chlorine and sodium ions were added to reach a concentration of approximately 150 mM and neutralize the total charge of the systems.

### 2.2. Molecular dynamics simulations with AmberTools and the Amber ff14SB force field

Molecular dynamics (MD) simulations of preconstructed SLN derived systems were performed using the Amber ff14SB force field^19^ and Lipid17^20^ topologies and parameters implemented in AmberTools 18 package^21^. We employed the TIP3P model to simulate water molecules. The systems were energy minimized and equilibrated following the six-step preparation protocol recommended by CHARMM-GUI. Temperature was maintained at 310 K with Langevin thermostat algorithm and pressure was set to 1.0 bar using the Monte Carlo (MC) barostat. All bonds involving hydrogens were constrained using the SHAKE algorithm. The systems were subjected to energy minimization followed by equilibration as follows: two rounds of 25-ps MD simulation using a canonical ensemble (NVT) and one 25-ps MD simulation using an isothermal-isobaric ensemble (NPT), and three independent 100-ps MD simulations using the NPT ensemble. We used the equilibrated systems as a starting point to perform five independent 1 μs MD replicates of each SLN truncation construct. Additionally, we carried out three independent 0.5 μs MD replicates of constructs SLNWT, SLN_1-26_, and SLN_1-22_ in a 1,2-dimyristoyl-sn-glycero-3-phosphocholine (DMPC) lipid bilayer following the same methodology.

### 2.3. Molecular dynamics simulations with NAMD and the CHARMM36 force field

We carried out six independent 1 μs MD replicates of constructs SLNWT, SLN_1-26_, and SLN_1-25_ using CHARMM36 force field^22^ implemented in NAMD software^23^. MD simulations were performed with periodic boundary conditions, particle mesh Ewald, and the RATTLE algorithm to constrain bonds to hydrogen atoms and allow a 2-fs time step. The NPT ensemble was maintained with a Langevin thermostat (310K) and an anisotropic Langevin piston barostat (1 atm). Solvated systems were first subjected to energy minimization, followed by gradually warming up of the systems for 200 ps. This procedure was followed by 20 ns of equilibration with backbone atoms harmonically restrained using a force constant of 10 kcal mol^−1^ Å^−2^. Production runs were continued for 1 μs each.

### 2.4. Analysis and visualization

The insertion of the TM in the membrane was obtained by computing the difference between the center of mass of both the TM backbone and the POPC bilayer. TM tilt angles relative to the membrane normal and dihedral angles were calculated using *helanal* and *dihedral* modules of the MDAnalysis python library^24^. Local membrane property analysis, per-residue contacts between SLN and the POPC headgroups, and secondary structure were computed with *g_lomepro*, *mindist*, and *do_dssp* tools implemented in GROMACS program, respectively.^25,26^ Mutual information analysis for the last 800 ns from the five MD replicates of SLN were performed with MutInf software^27^. Figures were generated with PyMOL v1.7^28^ and plots were constructed with Gnuplot v5.2^29^.

## 3. RESULTS AND DISCUSSION

### 3.1. CT deletions do not alter SLN insertion in the lipid bilayer

SLN_WT_ shows a behavior very similar to that observed in nuclear magnetic resonance (NMR) and MD studies carried out by other research groups.^30^ Root-mean-square deviation (RMSD) analysis showed that serial deletion of SLN’s CT does not affect the stability of the secondary structure of the protein in the μs time scale (**Figures S2–S11**). Root-mean-square fluctuation (RMSF) (**Figures S2–S11**) and secondary structure (**Figure S13**) analyses revealed that the TM domain of SLN is structurally stable in all SLN_WT_ and SLN serial deletion constructs. These analyses imply that deletions in the CT domain, including those beyond the CT domain (i.e., SLN_1-22_ through SLN_1-26_), do not alter the structural stability of SLN in the conditions used here.

After having established that SLN deletion constructs are structurally stable in a lipid environment, we tested the cell-based hypothesis that the CT domain is needed for retention in the lipid bilayer.^15^ Lipid insertion analysis showed that incorporation of all SLN deletion constructs in the membrane is similar to that of SLN_WT_ (**Figure 2A**), indicating that serial deletion of residues in the CT does not induce dissociation of SLN from the membrane. Nevertheless, we found that serial deletion of residues in the CT domain promotes a gradual decrease in the TM tilt angle (**Figure 2B**) and the local thickness of the lipid bilayer (**Figure 2C** and **Figure S12**).

**Figure 2.**
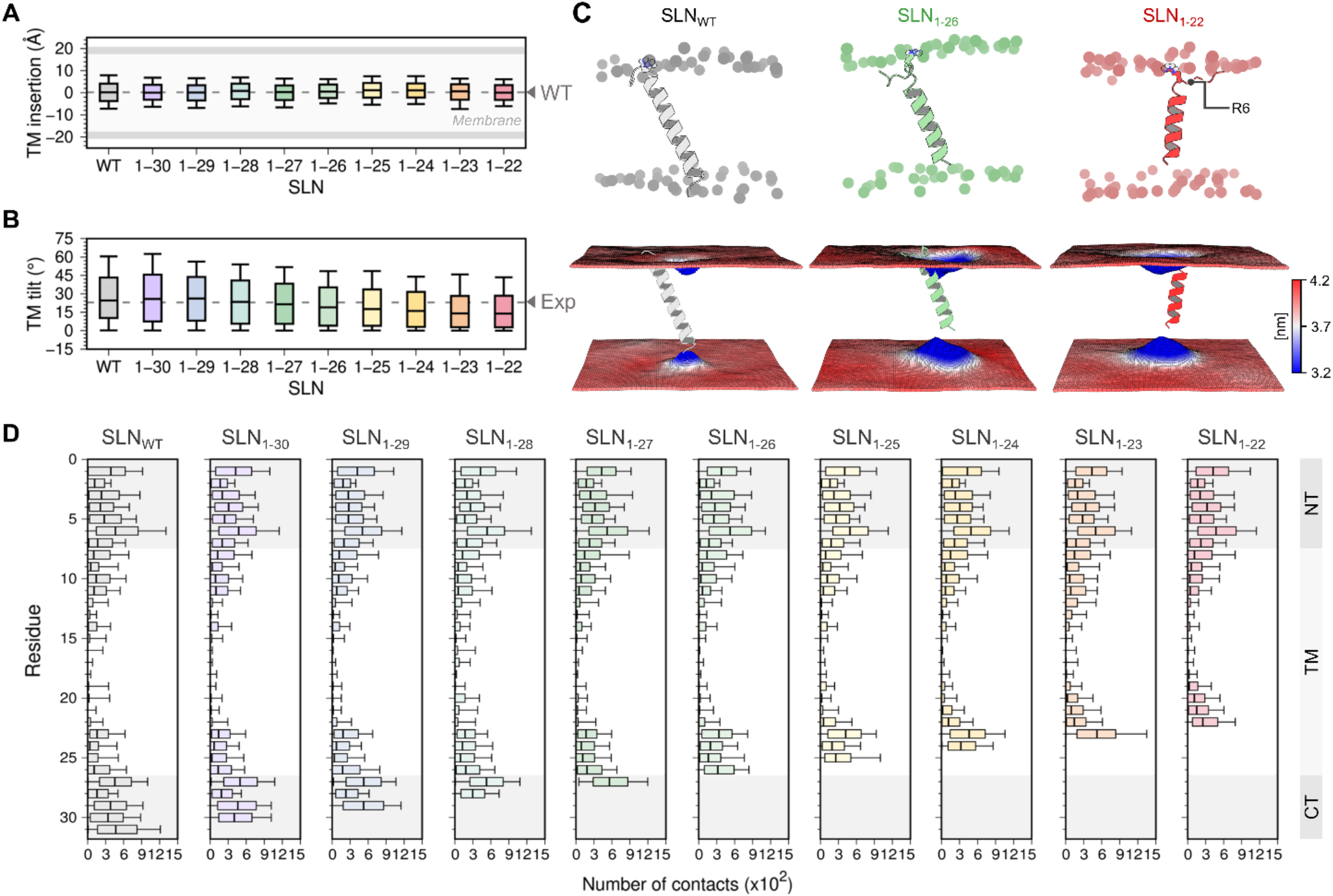
Orientation and lipid/protein interaction profiles of SLNWT and SLN deletion constructs. (A) Lipid insertion profile relative to the center of mass of both the protein and the lipid bilayer. (B) Tilt angle of the TM domain relative to the membrane normal. The average experimental tilt angle of the TM domain is 23° (Exp., dashed gray line).^10^ (C) Representative orientation (*top*) and time averaged local membrane thickness (*bottom*) of constructs SLNWT (gray, *left*), SLN_1-26_ (green, *center*), and SLN_1-22_ (red, *right*). Local membrane property analysis was performed with the *g_lomepro* package.^25^ (D) Number of contacts between side chain atoms and lipid headgroup using a cutoff distance of 6.0 Å; shaded areas show the location of NT and CT domains of SLN. All box plots show the full range of variation (from minimum to maximum) using 2.5 %, 50.0 %, and 97.5 % quantiles.

The average tilt angle computed for SLNWT in the MD trajectories reported here is 25°; this value is in remarkable agreement with average tilt angle values of 23° and 24° obtained using solid-state NMR (ssNMR)^10^ and hybrid magic angle spinning/oriented solid-state NMR spectroscopy (MAS/O-ssNMR)^31^, respectively. Furthermore, the tilt angle and structural stability of SLNWT are in good agreement with previous MD simulations in which different lipid bilayer models were used.^30,32,33^ Membrane thickness is an important variable that influences the behavior of membrane proteins.^34^ Indeed, Monte Carlo simulations carried out using a ~30 Å thick implicit membrane model showed that SLN has an average tilt angle of 16 ± 8° under these conditions.^32^ A 5° increase in the tilt angle was observed for SLN embedded in DOPC membranes.^30,33^ These studies suggest that the thickness of lipid bilayer directly influences the orientation of SLN in the membrane. Therefore, it could be argued that the behavior of SLN in this study is an artifact produced by the choice of POPC to simulate the protein in a membrane environment. However, our simulations most likely capture the behavior of SLN in a native environment because (1) the lipid-protein system used in this study is consistent with the composition of the lipids of the SR membrane, i.e., ~68% of PC lipids;^35^ (2) the acyl chain length of POPC is optimal for the function of proteins found in the SR (i.e., C16–C18);^36^ and (3) exchange of PC with PE does not appear to have an effect on the structure of other key proteins found in the SR (e.g., SERCA).^37^

The effects of deletions in the CT domain are intriguing because changes in tilt angle and bilayer induced by SLN localization and retention upon serial deletions in the CT domain,^15^ but we do not observe substantial changes in the insertion profile of SLN deletion constructs (**Figure 2A**). This apparent contradiction can be reconciled by analyzing the contacts between each residue of SLN and the lipid headgroups (**Figure 2D**). We focus on residues that play a key role in anchoring and stabilization of proteins in the membrane, such as arginine^39^ and tryptophan.^40^ We found that in SLNWT as well as SLN deletion constructs SLN_1-27_ through SLN_1-30_, anchoring residues R6 and R27 (located at the NT and CT domains, respectively) exhibit the highest number of contacts with the lipid headpiece. Serial deletion of CT residues beyond R27 does not induce loss of protein/lipid headpiece contacts involving the NT domain, and the interactions involving anchoring residue R6 remain intact even in the shortest SLN deletion construct SLN_1-22_ (**Figure 2D**). In the absence of residue R27 in CT domain, we also observed a substantial increase in the number of contacts between residue W23 and the lipid headgroups (**Figure 2D**), thus indicating that W23 serves as a surrogate anchor in the absence of R27. The latter agrees with MAS/O-ssNMR data indicating that R6, W23 and R27 play a central role in anchoring SLN to the membrane and modulating its tilt angle.^31^ These findings suggest that while CT residues play a role in SLN preserving normal orientation of SLN in the membrane, the CT domain is not absolutely required for retention and localization. Preservation of the TM domain topology and the presence of native and surrogate anchor residues upon serial deletions in the CT domain help explain experimental evidence showing that SLN deletion constructs are able to deletions are consistent with adaptation to hydrophobic mismatch.^38^ This effect correlates with impaired membrane localize to the membrane when they are co-expressed with cardiac SERCA.^15^

### 3.2. CT domain controls interdomain communication in SLN

Closer inspection of the distribution of ϕ/ψ torsion angles in the NT domain led to the interesting observation that serial deletions in the CT induces changes in the backbone dynamics of cytosolic residues I3-E7 (**Figure S14**). The distribution of ϕ/ ψ torsion angles indicates that serial deletion of CT residues shifts the equilibrium away from the right-handed α-helix region populated in the SLN_WT_ toward the β-sheet and poly-Pro II regions of the ϕ/ψ dihedral angle space (**Figure S14**). The shift in the ϕ/ψ torsion angles induced by serial deletions in SLN implies that the CT domain exerts control over the structural dynamics of the NT domain. To test this hypothesis, we performed mutual information analysis^27^ of the dihedral distributions to identify correlated motions between the NT and CT domains of SLN. We found that in SLN_WT_, there is a strong correlation between residues G2-L8 and V26-Q30 (**Figure 3**). Correlated motions between the CT and NT domains are also observed in SLN_1-30_, and SLN_1-29_ and to some extent in SLN_1-28_; however, serial deletions beyond residue S28 partially or completely abolish the ability of the CT domain to communicate with the cytosolic NT domain of SLN (**Figure 3**). This mechanistic evidence indicates that the CT segment dynamically controls interdomain communication in SLN, and correlates well with experiments showing that deletion of conserved residues R27–Y31 results in loss of function.^14,15^

**Figure 3.**
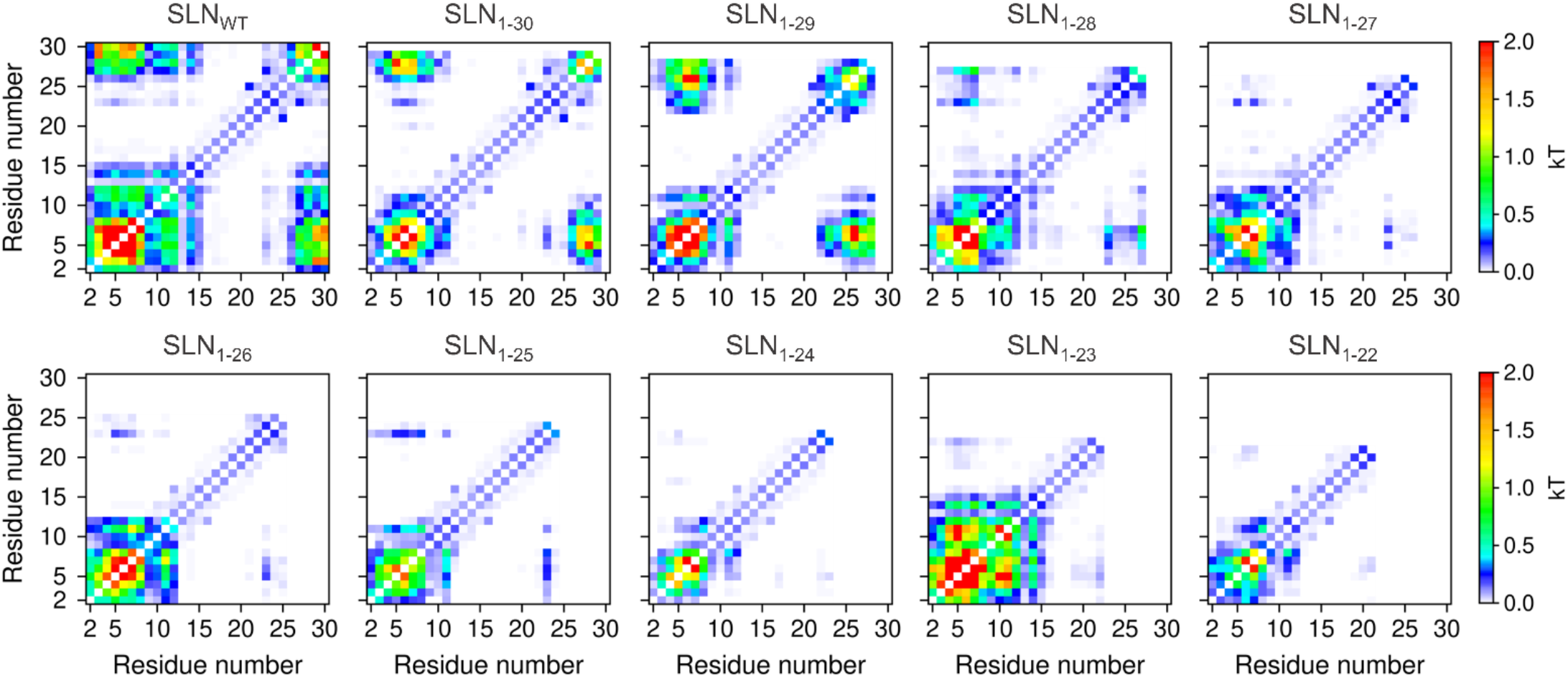
Mutual information matrix between residues calculated from the trajectories of SLN_WT_ and the nine serial deletion constructs of SLN. Mutual information analysis describes correlations between torsional degrees of freedom from an expansion of the molecular configurational entropy, and is expressed in units of kT.^27^ Analysis was performed using the MutInf software package.^27^

Intrinsic ϕ/ψ distributions for individual amino acids which are known to be force-field dependent,^41^ so it is possible that the correlated motions between the NT and CT domains is an artifact of the force field and MD engine used in this study. To rule out this possibility, we performed additional simulations of SLN_WT_, SLN_1-26_, and SLN_1-25_ employing the CHARMM36 force field^22^ and the NAMD software^23^. We found that SLN constructs simulated with the CHARMM36 force field are structurally stable in the μs timescale, do not dissociate from the membrane upon deletion of the CT domain, and reproduce the decrease in tilt angle and the loss of correlated motions between the NT and CT observed in the simulations performed with the Amber force field (**Figures S15–S18**). A similar behavior is observed in MD simulations of constructs SLN_WT_, SLN_1-26_, and SLN_1-22_ with the Amber force field using a thinner membrane containing DMPC lipids (**Figures S19–S22**). Due to the nature of the lipid bilayer, we found that the tilt angle of SLNWT changes from 25.0° in a POPC bilayer to a value of 35.1° in the DMPC bilayer; similar changes were also observed in the constructs SLN_1-26_, and SLN_1-22_ (**Figure S20**). Despite these changes in tilt angle, the constructs embedded in a DMPC lipid bilayer are structurally stable and do not dissociate from the membrane upon deletion of the CT domain. More importantly, we found that the decrease in tilt angle and the loss of correlated motions induced by deletion of the CT domain are independent of lipid composition. These findings, which are highly reproducible across force fields and membrane thickness, demonstrate that the interdomain communication between the NT and CT domains is an intrinsic functional feature of SLN.

### 3.3. Phosphorylation of residue T5 induces changes in SLN interdomain communication

We also asked whether phosphorylation of residue T5, a posttranslational modification that regulates SLN function in the cell,^42^ affects interdomain communication in SLN. We performed atomistic μs-long MD simulations of SLN phosphorylated at residue T5 (SLN_pT5_). SLNpT5 is structurally stable in the μs timescale (**Figure S23**), and that phosphorylation of SLN does not induce substantial changes in membrane insertion (**Figure 4A**) or tilt of the TM domain (**Figure 4B**). However, phosphorylation of T5 induces a shift in the ϕ/ψ torsion angles toward the β-sheet regions of the dihedral angle space (**Figure 4C**), and also induces a decrease in the correlation between NT and CT domains (**Figure 4D**). These findings further support our MD-based conclusion that there is dynamic communication between the CT and NT domains, and indicates that the phosphorylation site at residue T5 serves as a switch that modulates interdomain communication to regulate SLN-mediated regulation of SERCA.^42^ The latter may explain the loss of SLN inhibitory activity exerted over SERCA when the peptide is phosphorylated, suggesting that communication between domains contributes to signal integration underlying SLN-mediated regulation of SERCA.^42^ This mechanistic evidence also supports the notion that the CT and NT are functionally distinct, yet essential domains in the regulation of SERCA.^11,14,43^

**Figure 4.**
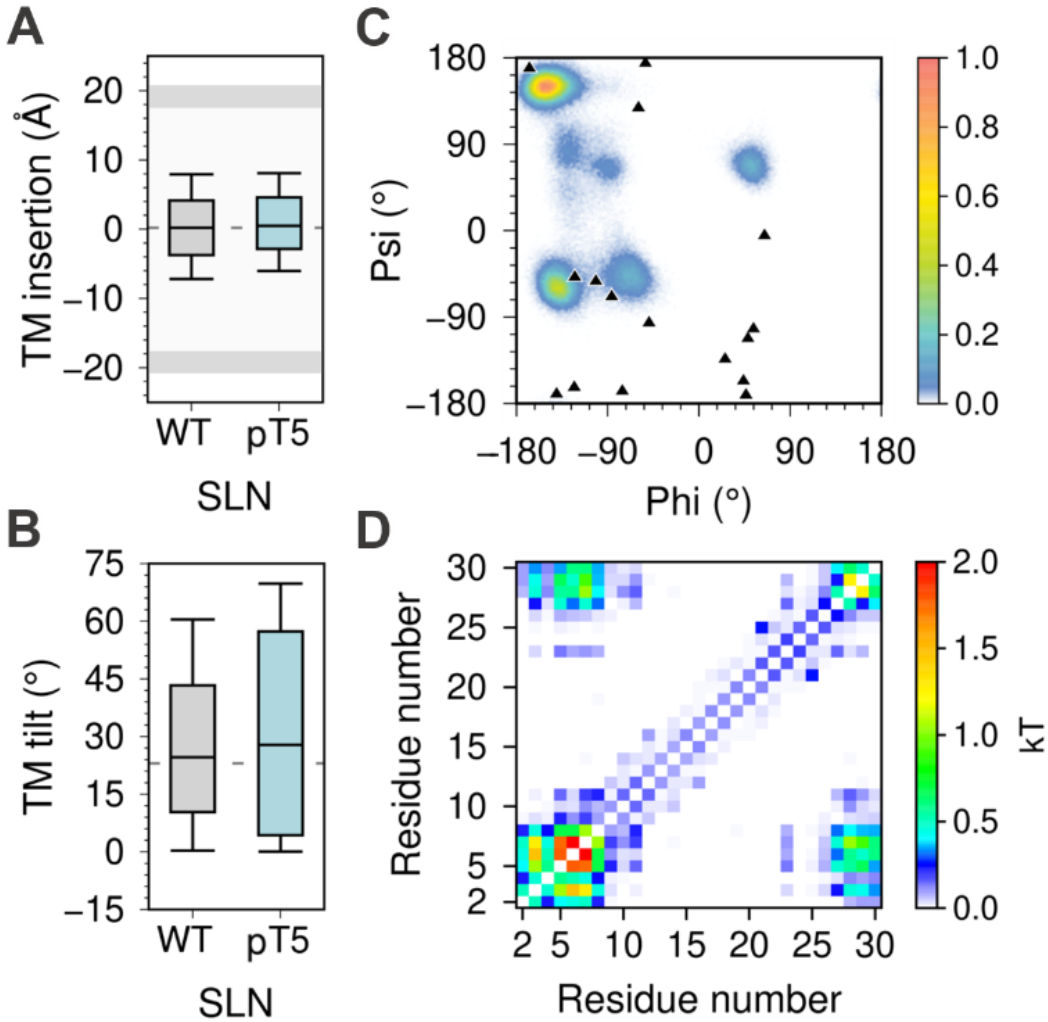
MD simulations analysis of phosphorylated SLN. (A) membrane insertion and (B) tilt angle of the TM domain in the lipid bilayer. (C) Dihedral angle distribution of residue T5; the ϕ/ψ torsion angles calculated from the 16 NMR models of SLN (PDB: 1JDM^9^) are shown as black triangles. All box plots show the full range of variation (from minimum to maximum) using 2.5 %, 50.0 %, and 97.5 % quantiles. (D) Mutual information matrix between residues calculated from the trajectories of phosphorylated SLN.

### 3.4. Potential functional roles of interdomain communication in SLN

The correlated motions between the NT and CT segments in SLN suggests that interdomain communication may play a role in the functional outcomes of the protein. We propose two hypotheses on the functional role of interdomain communication in the context of SERCA regulation: (1) The CT segment may exert dynamic control over the NT segment to regulate SERCA through a similar mechanism used by PLB, where order-to-disorder transitions of the NT segment dictate inhibition or activation of the pump.^44^ (2) X-ray crystallography studies^12^ have shown that the NT domain of SLN interacts with SERCA via bridge-mediated interactions, and MD simulations have shown that such interactions induce a structural rearrangement in the energy-transduction domain of SERCA, thus perturbing Ca^2+^ occlusion in the transport sites and contributing to Ca^2+^ backflux to the cytosol.^43^ Therefore, it is possible that the correlated movements within SLN may control the interactions between SERCA and the NT segment of SLN, thereby modulating uncoupling of SERCA’s Ca^2+^ transport and ATP hydrolysis.

## 4. CONCLUSION

In summary, we used extensive atomistic MD simulations to elucidate the function role of the conserved CT domain of SLN by studying the effect of nine serial deletions in this domain. We find that wild-type SLN and deletion constructs are structurally stable and do not dissociate from the membrane in the μs time scale. However, we found that interdomain communication between the CT and NT domains intrinsic to SLN is abolished upon deletion the conserved segment Tyr29-Tyr31 as well as by serial deletions beyond this domain. This evidence suggests that the CT domain dynamically controls interdomain communication in SLN. This mechanistic evidence is further supported by MD simulations of SLN showing that phosphorylation of residue T5 induces a decrease in the correlation between NT and CT domains. Based on this extensive evidence, we propose the novel hypothesis that the conserved C-terminal domain is an essential element required for dynamic control of SLN regulatory function. Our study also provides structural hypotheses that can be tested experimentally. For example, EPR spectroscopy using the backbone spin label TOAC can be used to measure with high precision changes in structural dynamics of the NT upon deletion of CT residues and SLN phosphorylation.^45,46^ These findings have important implications for understanding the molecular mechanisms for interdomain communication in small regulatory proteins, and how this shapes their function in the cell.

## ASSOCIATED CONTENT

Supporting Information

The Supporting Information is available free of charge Figure S1: Multiple alignment of SLN mammalian amino acid sequences. Figures S2-S11: Backbone RMSD and per-residue RMSF of SLN deletion constructs. Figure S12: Helical content of SLN deletion constructs. Figure S13: TM domain orientation and time-averaged local membrane thickness of SLN deletion constructs. Figure S14: Distributions of the φ/ψ backbone dihedral angles of SLN deletion constructs. Figures S15-S18: Backbone RMSD, RMSF, TM insertion, tilt angle, φ/ψ torsion angle distribution, mutual correlation analysis, helical content, and contacts with the lipid headgroups of three SLN deletion constructs simulated with the CHARMM36 force field. Figures S19-S22: Backbone RMSD, RMSF, TM insertion, tilt angle, φ/ψ torsion angle distribution, mutual correlation analysis, helical content, contacts with the lipid headgroups, and membrane thickness analysis of three SLN deletion constructs simulated with the Amber ff14SB force field in the DMPC lipid bilayer. Figure S23: Backbone RMSD and per-residue RMSF of SLN phosphorylated at residue T5.

## AUTHOR INFORMATION

### Notes

The authors declare no competing financial interest.

## ACKNOWLEDGMENTS

This work was supported by the National Institutes of Health grant R01GM120142. This research was supported through computational resources and services provided by Advanced Research Computing at the University of Michigan, Ann Arbor.

