## Supplementary Information for "Conserved luminal C-terminal domain dynamically controls interdomain communication in sarcolipin"

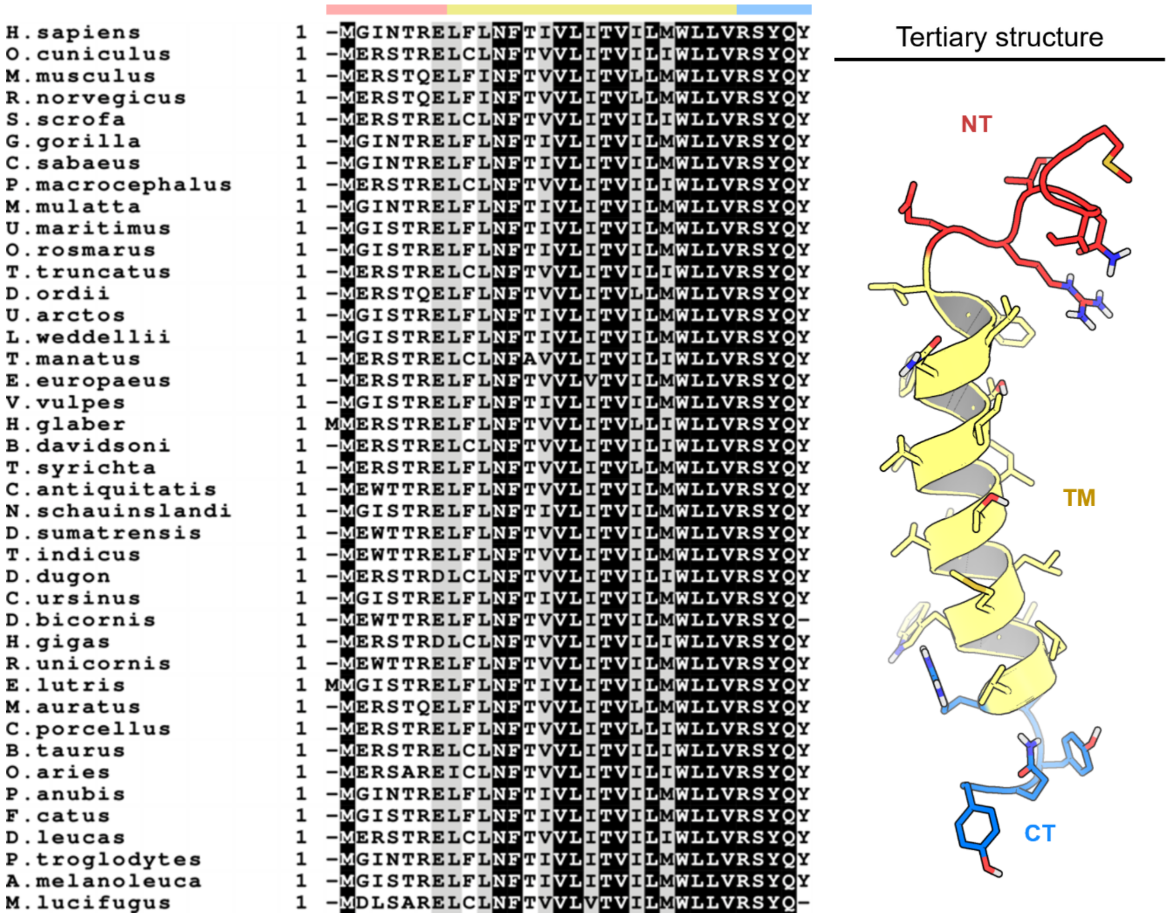


**Figure S1.** Multiple alignment of mammalian amino acid sequences and depiction of the tertiary structure of SLN. Sequence alignment was carried out with T-Coffee web server[1]. SLN is shown as ribbon and stick representation and colored according to its functional domains: NT (purple), TM (gray), and CT (orange).

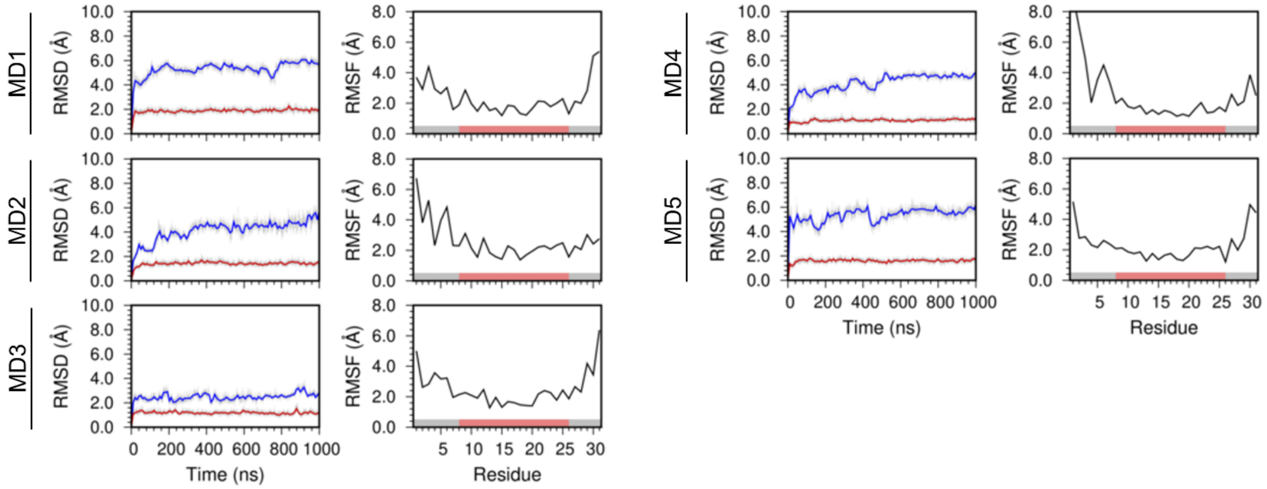


**Figure S2.** Backbone RMSD and per-residue RMSF of $SLN_{WT}$. The RMSD of the full-length protein (blue) and the TM domain (red) was calculated by aligning the backbone of the TM domain with the structure at the beginning of each 1-µs MD replicate simulated with the Amber ff14SB force field. RMSFs of Cα atoms were calculated from each independent MD trajectory using the backbone of the TM domain as a reference.

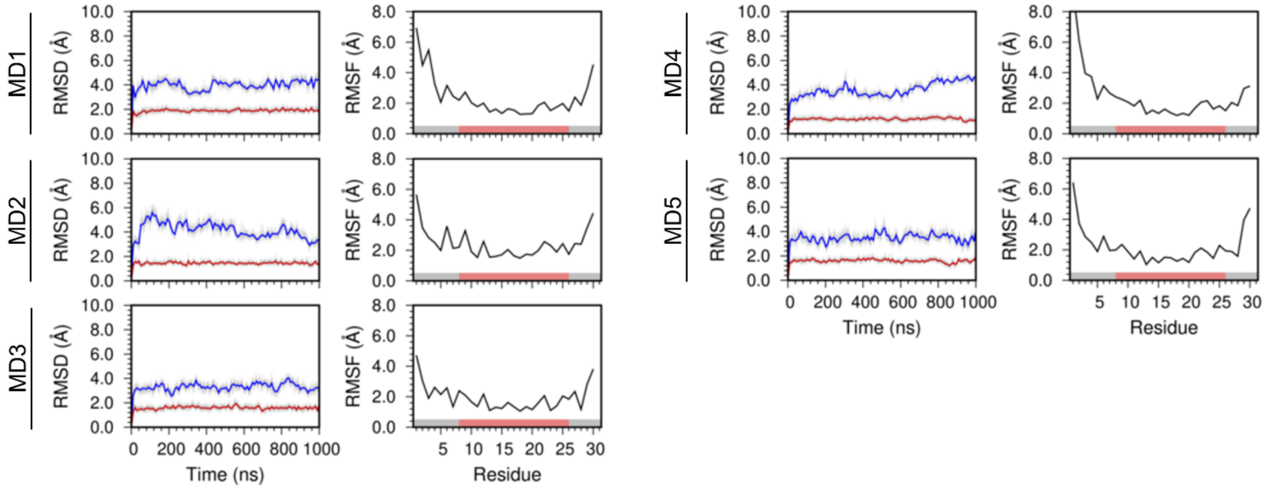


**Figure S3.** Backbone RMSD and per-residue RMSF of SLN$_{1\text{-}30}$. The RMSD of the full-length protein (blue) and the TM domain (red) was calculated by aligning the backbone of the TM domain with the structure at the beginning of each 1-μs MD replicate simulated with the Amber ff14SB force field. RMSFs of Cα atoms were calculated from each independent MD trajectory using the backbone of the TM domain as a reference.

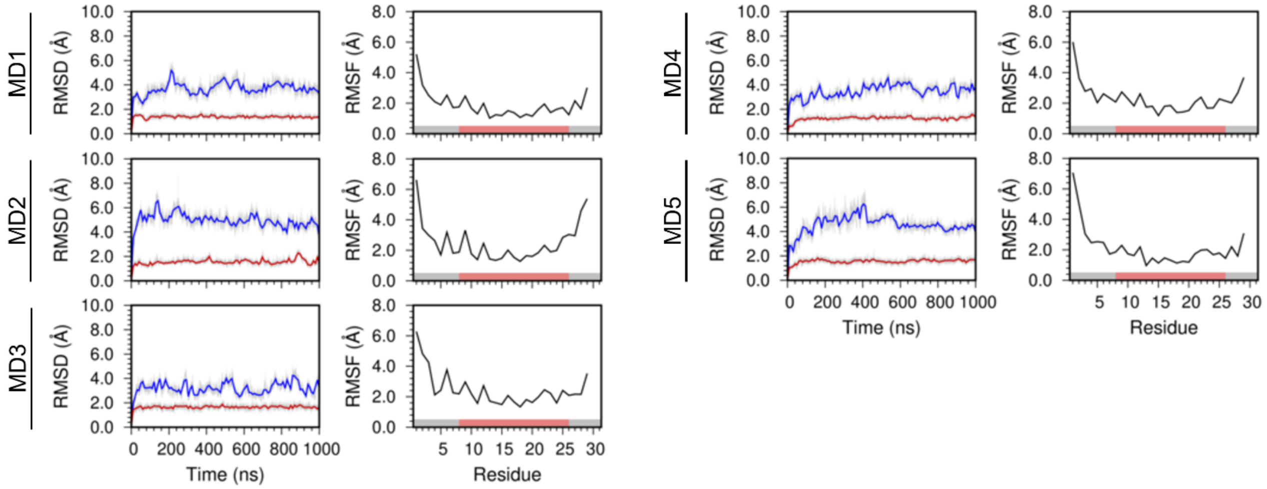


**Figure S4.** Backbone RMSD and per-residue RMSF of $SLN_{1\text{-}29}$. The RMSD of the full-length protein (blue) and the TM domain (red) was calculated by aligning the backbone of the TM domain with the structure at the beginning of each 1-µs MD replicate simulated with the Amber ff14SB force field. RMSF of Cα atoms were calculated from each independent MD trajectory using the backbone of the TM domain as a reference.

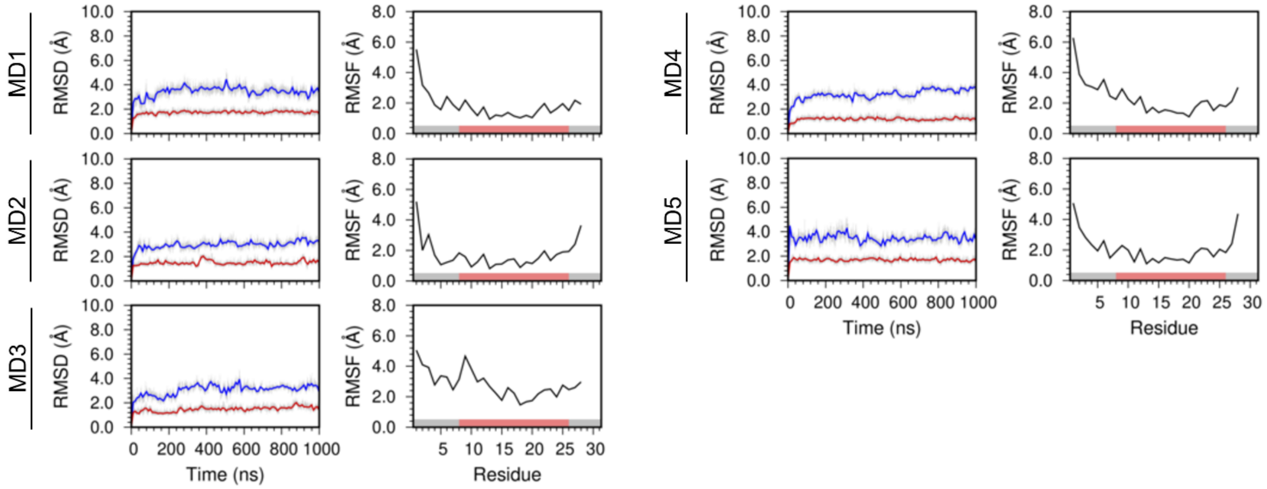


**Figure S5.** Backbone RMSD and per-residue RMSF of $SLN_{1\text{-}28}$. The RMSD of the full-length protein (blue) and the TM domain (red) was calculated by aligning the backbone of the TM domain with the structure at the beginning of each 1-μs MD replicate simulated with the Amber ff14SB force field. RMSF of Cα atoms were calculated from each independent MD trajectory using the backbone of the TM domain as a reference.

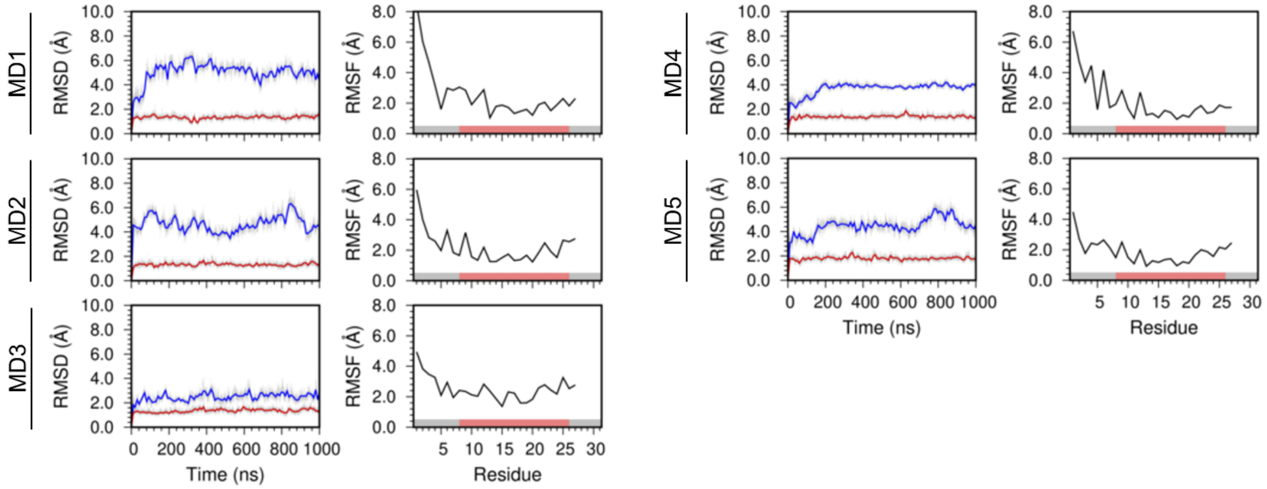


**Figure S6.** Backbone RMSD and per-residue RMSF of SLN1-27. The RMSD of the full-length protein (blue) and the TM domain (red) was calculated by aligning the backbone of the TM domain with the structure at the beginning of each 1-µs MD replicate simulated with the Amber ff14SB force field. RMSF of Cα atoms were calculated from each independent MD trajectory using the backbone of the TM domain as a reference.

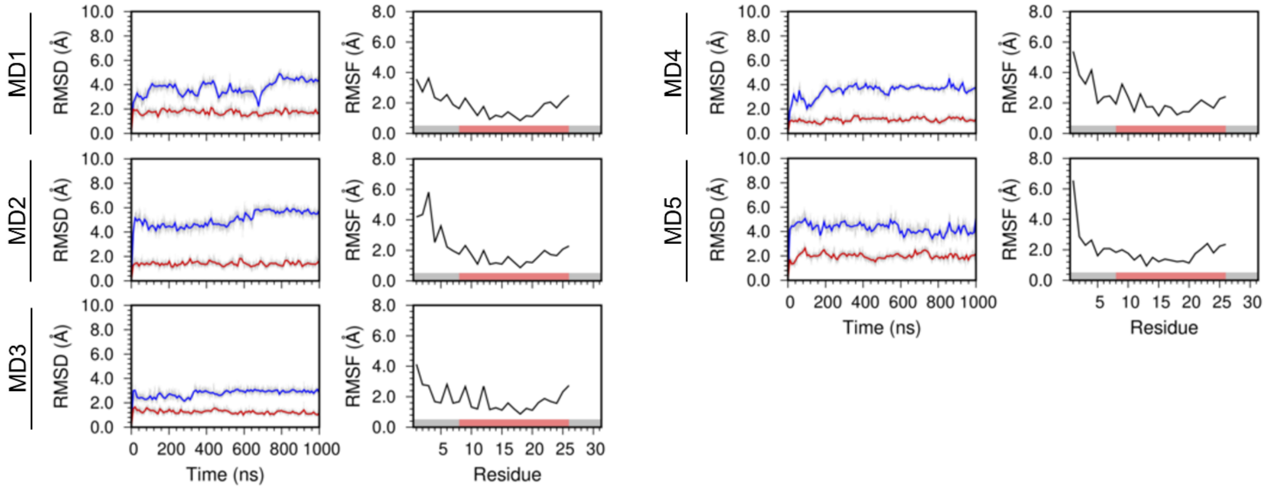


**Figure S7.** Backbone RMSD and per-residue RMSF of $SLN_{1\text{-}26}$. The RMSD of the full-length protein (blue) and the TM domain (red) was calculated by aligning the backbone of the TM domain with the structure at the beginning of each 1-µs MD replicate simulated with the Amber ff14SB force field. RMSF of Cα atoms were calculated from each independent MD trajectory using the backbone of the TM domain as a reference.

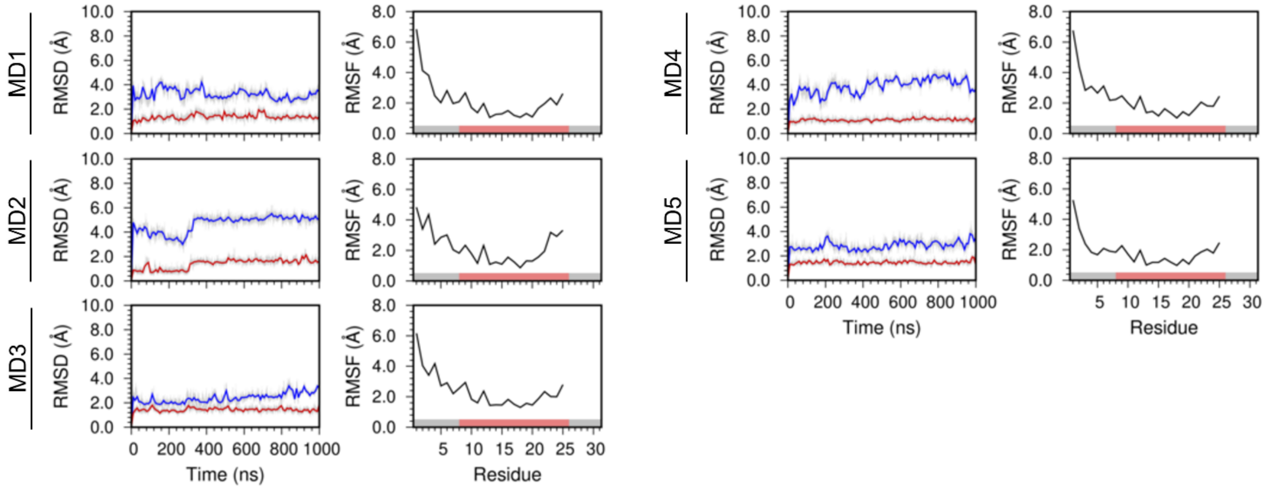


**Figure S8.** Backbone RMSD and per-residue RMSF of $SLN_{1\text{-}25}$. The RMSD of the full-length protein (blue) and the TM domain (red) was calculated by aligning the backbone of the TM domain with the structure at the beginning of each 1-µs MD replicate simulated with the Amber ff14SB force field. RMSF of Cα atoms were calculated from each independent MD trajectory using the backbone of the TM domain as a reference.

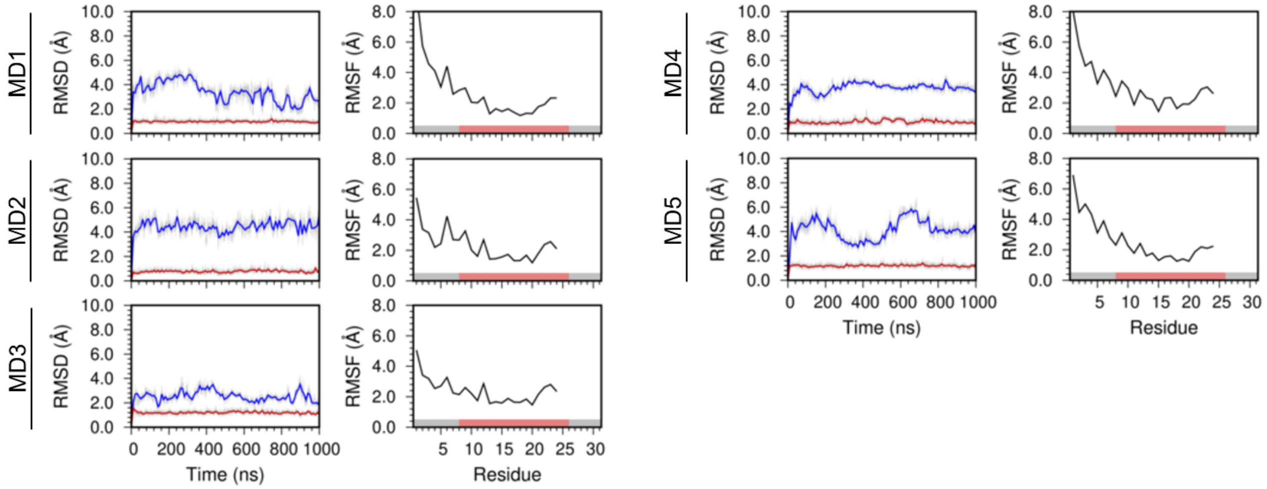


**Figure S9.** Backbone RMSD and per-residue RMSF of $SLN_{1\text{-}24}$. The RMSD of the full-length protein (blue) and the TM domain (red) was calculated by aligning the backbone of the TM domain with the structure at the beginning of each 1-µs MD replicate simulated with the Amber ff14SB force field. RMSF of Cα atoms were calculated from each independent MD trajectory using the backbone of the TM domain as a reference.

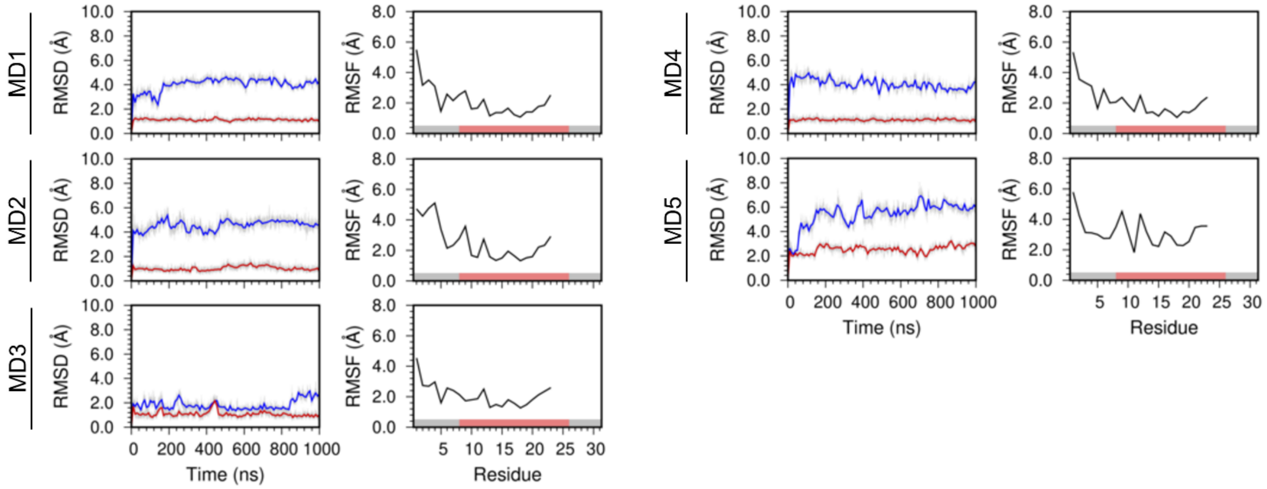


**Figure S10.** Backbone RMSD and per-residue RMSF of $SLN_{1-23}$. The RMSD of the full-length protein (blue) and the TM domain (red) was calculated by aligning the backbone of the TM domain with the structure at the beginning of each 1-µs MD replicate simulated with the Amber ff14SB force field. RMSF of Cα atoms were calculated from each independent MD trajectory using the backbone of the TM domain as a reference.

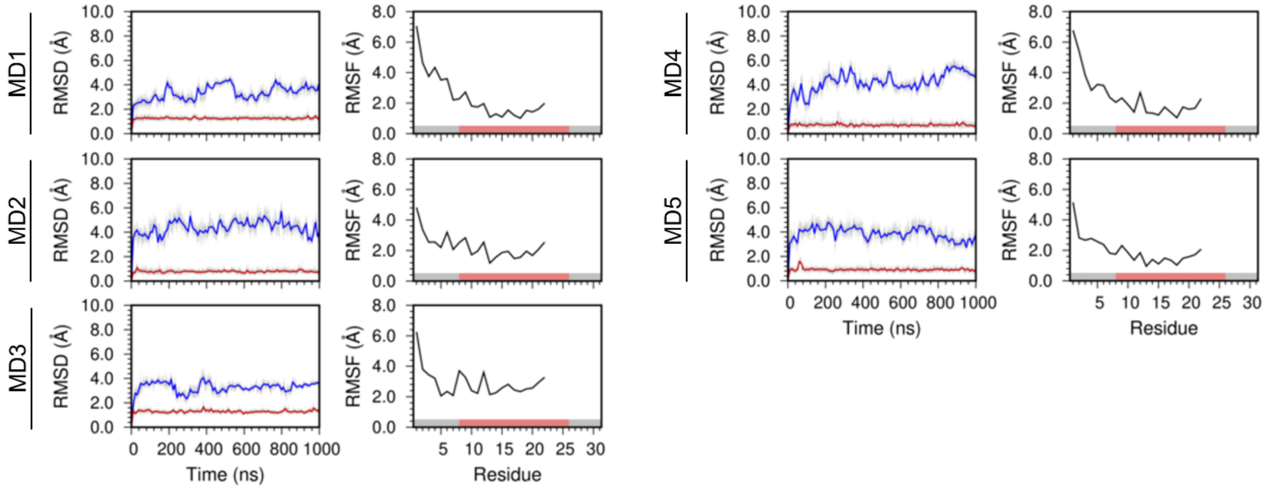


**Figure S11.** Backbone RMSD and per-residue RMSF of $SLN_{1\text{-}22}$. The RMSD of the full-length protein (blue) and the TM domain (red) was calculated by aligning the backbone of the TM domain with the structure at the beginning of each 1-µs MD replicate simulated with the Amber ff14SB force field. RMSF of Cα atoms were calculated from each independent MD trajectory using the backbone of the TM domain as a reference.

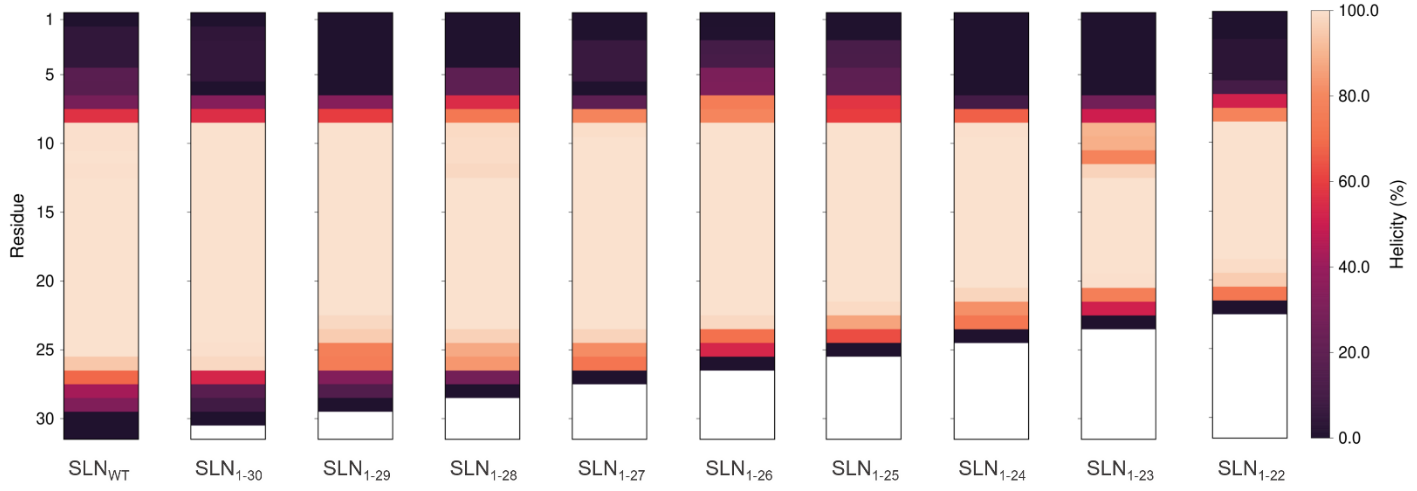


**Figure S12.** Helical content of SLN$_{WT}$ and SLN deletion constructs. The helical content was averaged over the five 1-µs MD replicates simulated with the Amber ff14SB force field; values are reported as % of helical content.

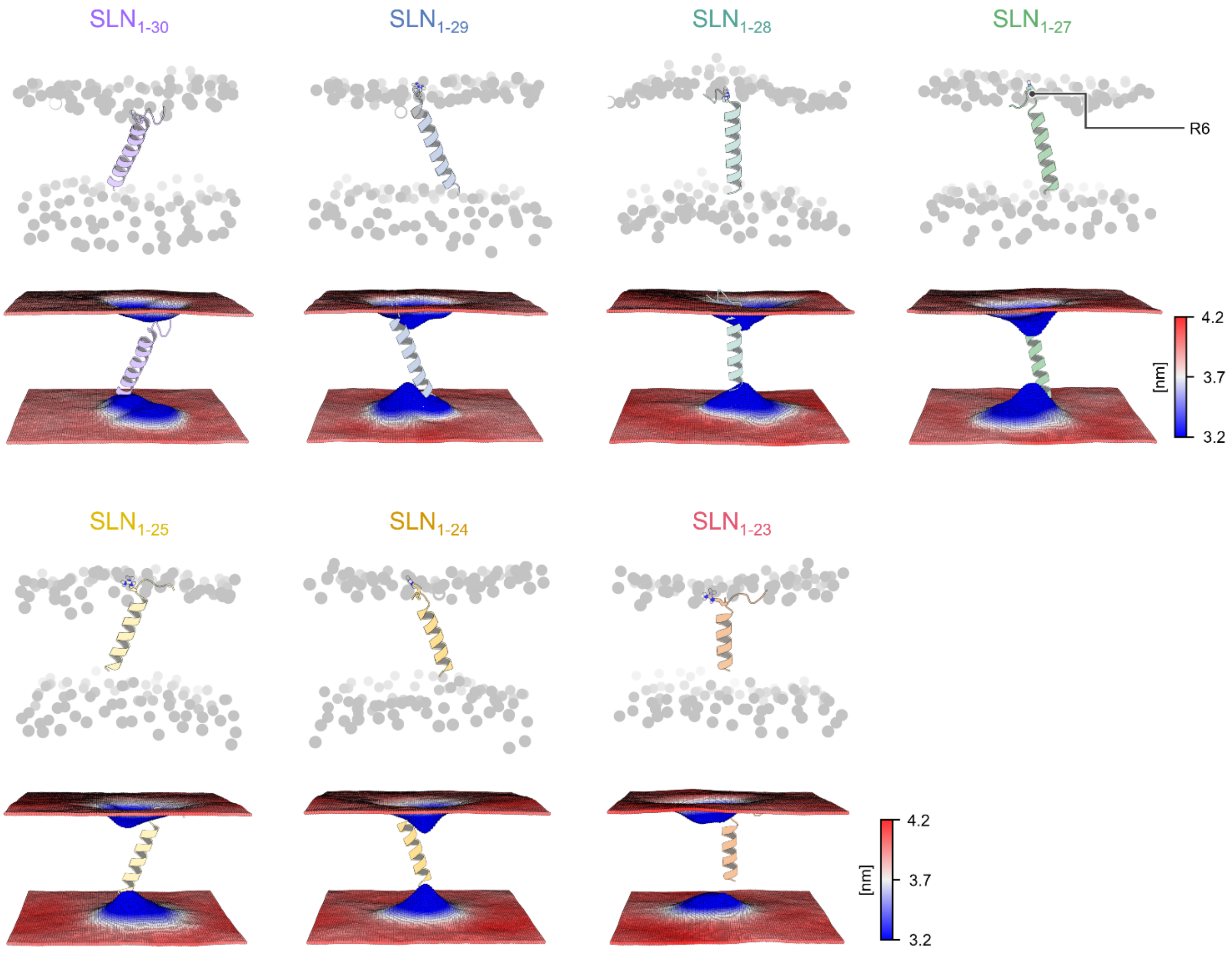


**Figure S13.** TM domain orientation and time-averaged local membrane thickness of the SLN deletion constructs. Representative structures of SLN constructs simulated with the Amber ff14SB force field (top) are shown as ribbons, and the lipid heads are shown as spheres. Local membrane property analysis (bottom) was performed with the *g_lomepro* package.[2]

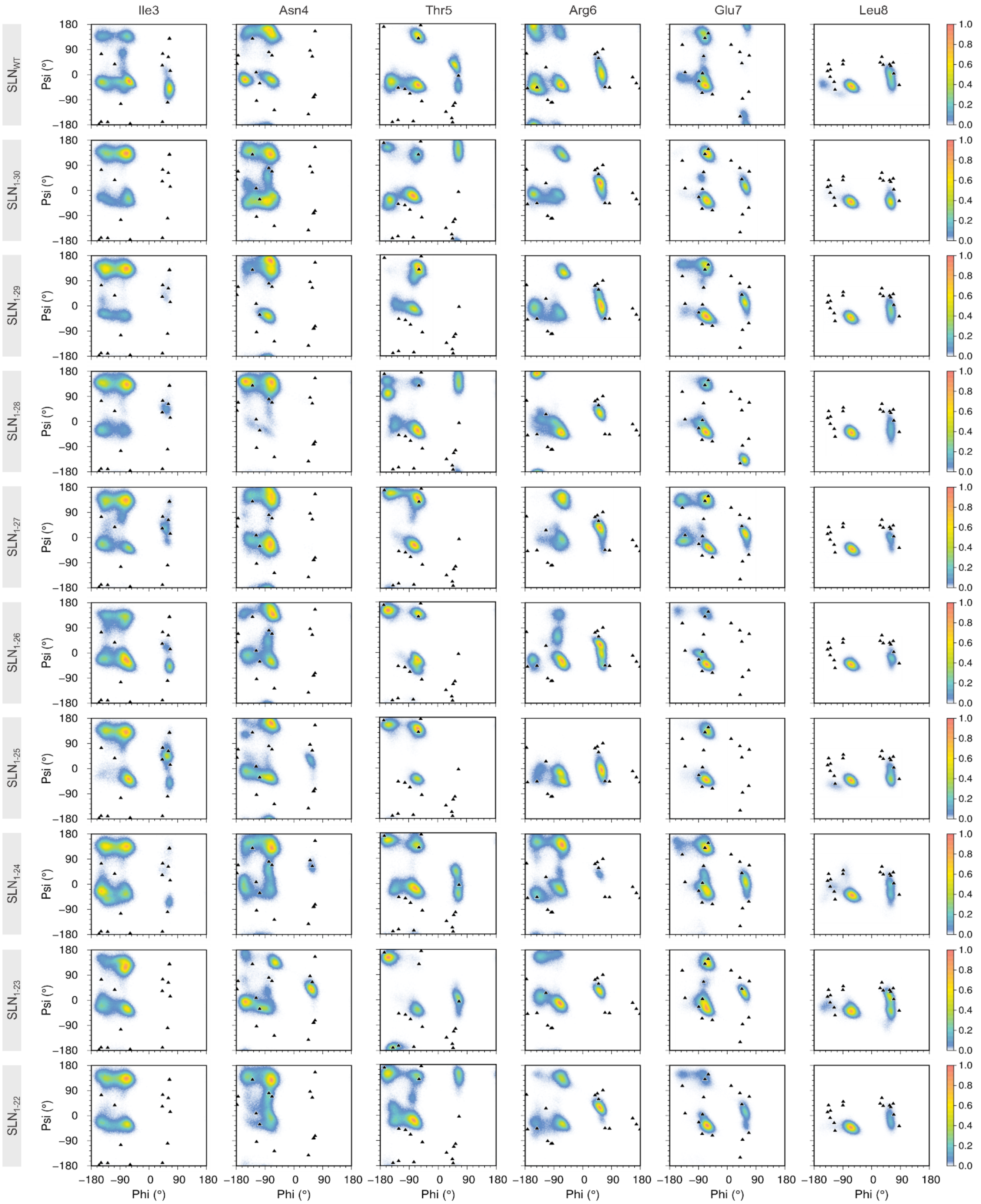


**Figure S14.** Distributions of the phi (φ) and psi (ψ) backbone dihedral angles of the NT domain amino acids of SLN$_{WT}$ and SLN deletion constructs. φ and ψ backbone dihedral angles were calculated over the combined five 1-μs MD replicates for each SLN construct simulated with the Amber ff14SB force field. For comparison, we show the φ/ψ dihedral angles calculated from each of the sixteen NMR models of SLN (black triangles, PDB: 1JDM).

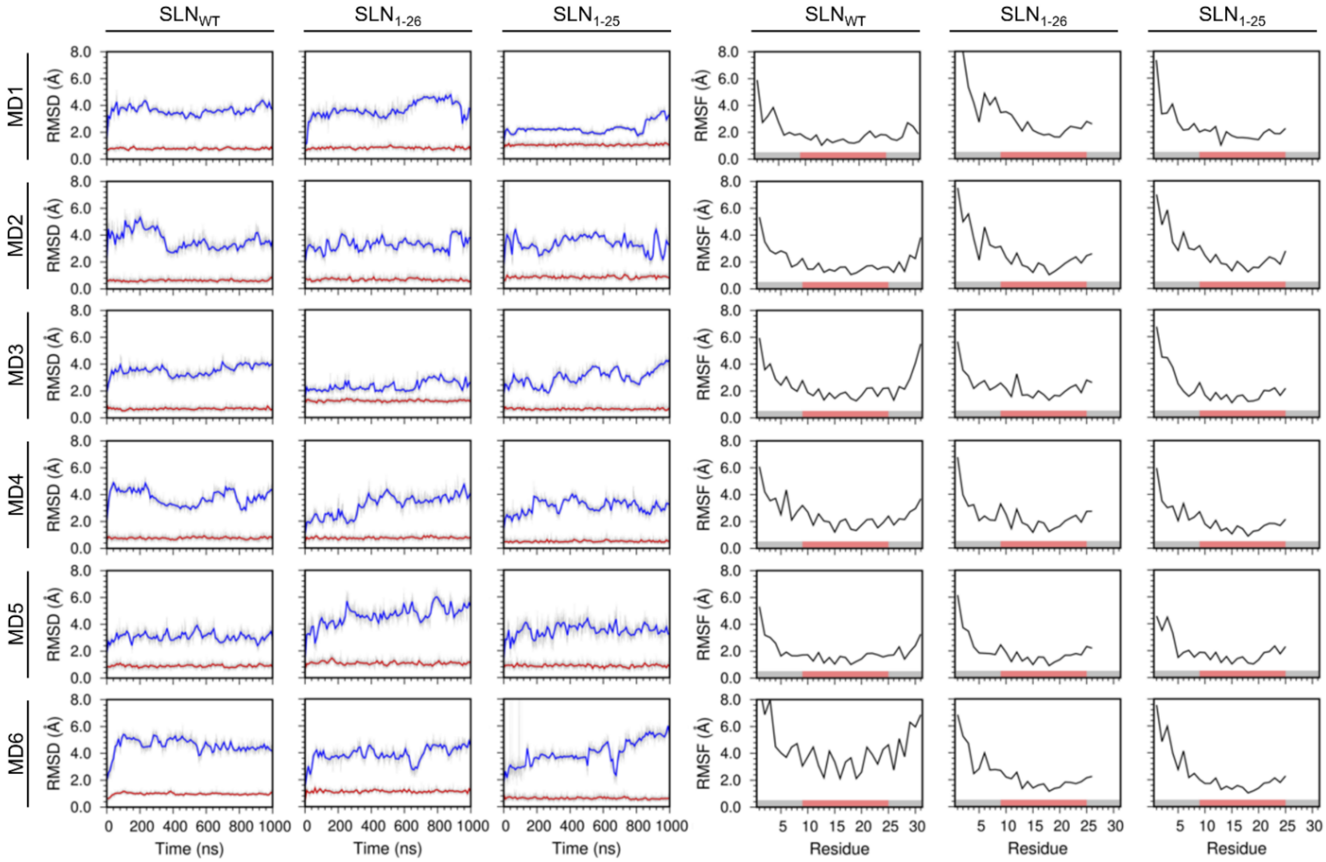


**Figure S15.** Backbone RMSD and per-residue RMSF of $SLN_{WT}$, $SLN_{1-26}$ and $SLN_{1-25}$ simulated with the CHARMM36 force field. The RMSD of the full-length protein (blue) and the TM domain (red) was calculated by aligning the backbone of the TM domain with the structure at the beginning of each 1-μs MD replicate. RMSF of Cα atoms were calculated from each independent MD trajectory using the backbone of the TM domain as a reference.

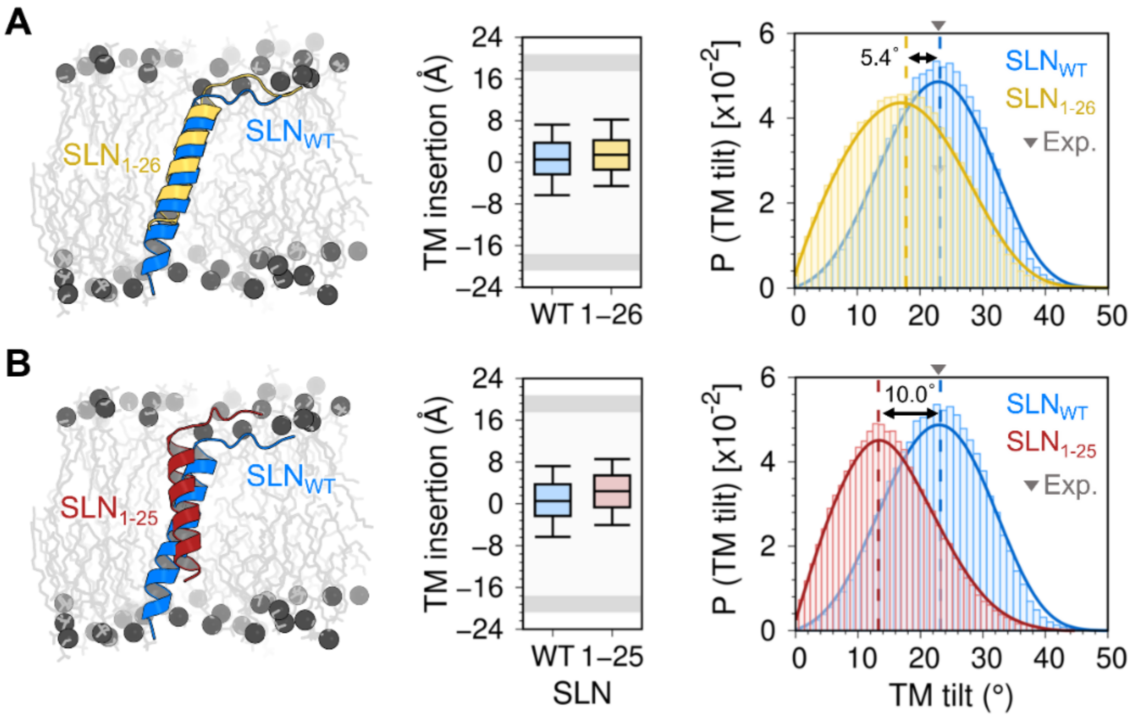


**Figure S16.** Comparison of the structure, TM insertion and tilt angle between SLN$_{WT}$ and (A) SLN$_{1\text{-}26}$ and (B) SLN$_{1\text{-}25}$ simulated with the CHARMM36 force field. For comparison, we superposed the most representative structures of SLN$_{WT}$ (blue ribbons) and the SLN deletion constructs (yellow and red ribbons); the lipid head and tails are show as gray spheres and sticks, respectively. Analysis of TM insertion and tilt angle was performed using all six MD trajectories combined; box plots show the full range of variation (from minimum to maximum) using 2.5 %, 50.0 %, and 97.5 % quantiles.

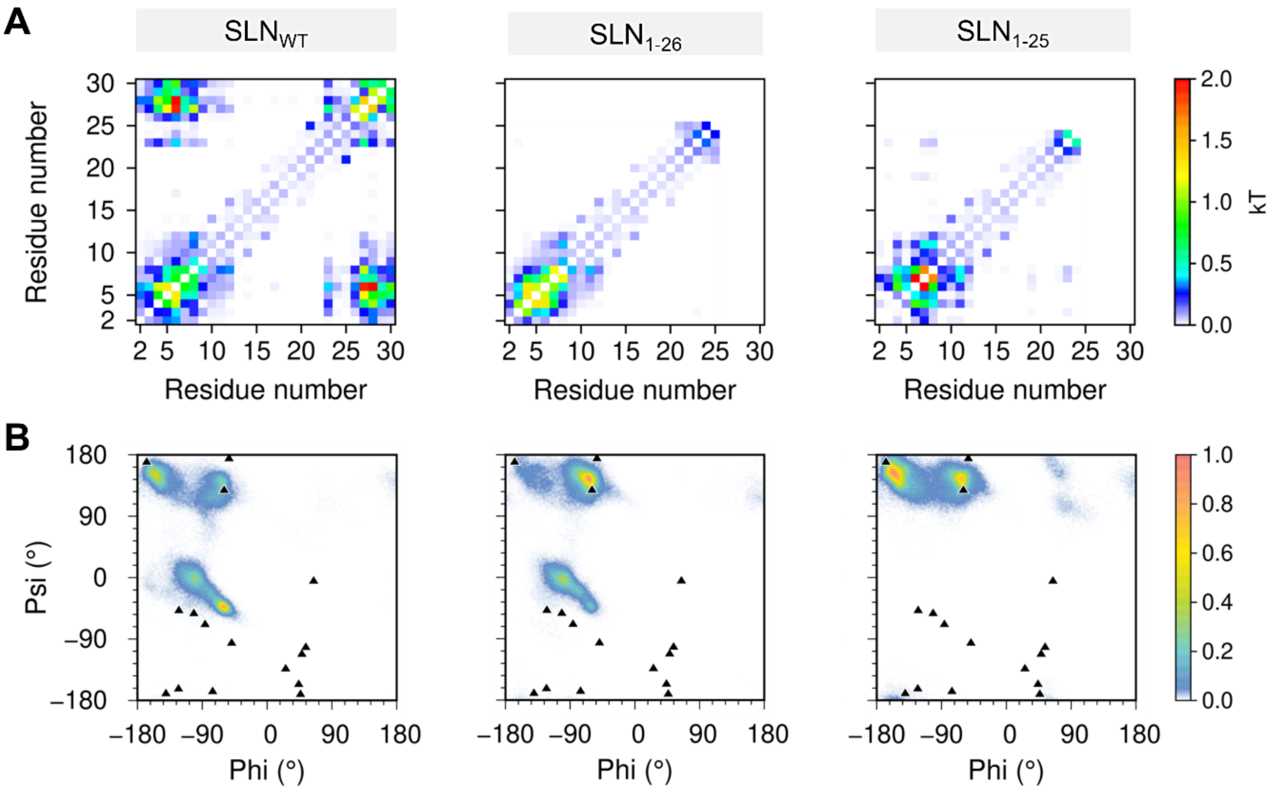


**Figure S17.** Mutual correlation analysis and φ/ψ torsion angle distribution of residue T5 obtained for $SLN_{WT}$ and constructs $SLN_{1-26}$ and $SLN_{1-25}$ simulated with the CHARMM36 force field. (A) Mutual information matrix between residues calculated from the trajectories of SLN constructs. (B) Dihedral angle distribution of residue T5, the phosphorylation site of SLN; the φ/ψ torsion angles calculated from the 16 NMR models of SLN (PDB: 1JDM[3]) are shown as black triangles.

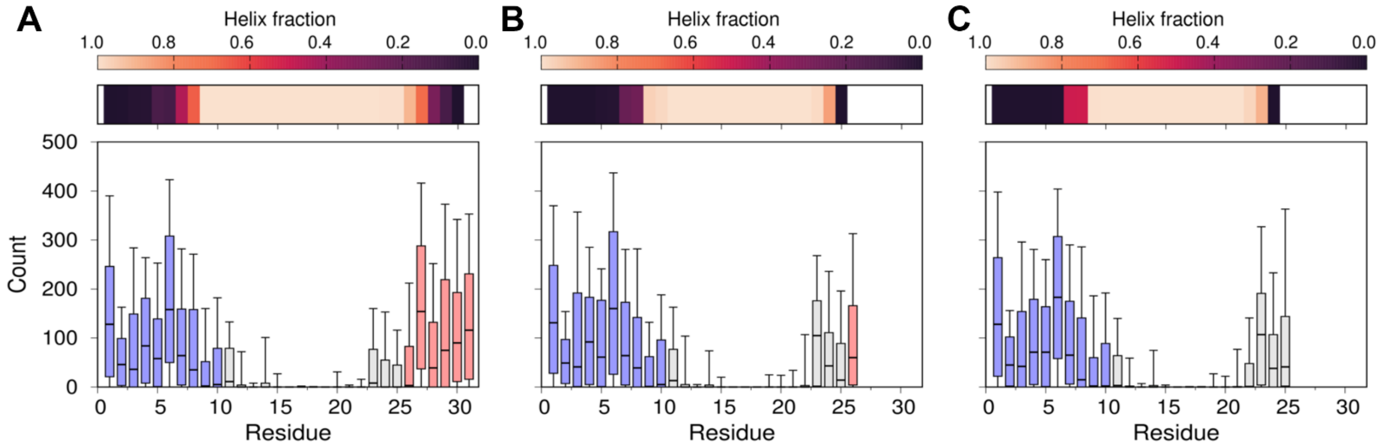


**Figure S18.** Helical content and number of contacts between side chain atoms and lipid headgroup calculated from the SLN constructs simulated with the CHARMM36 force field. Constructs simulated here are (A) $SLN_{WT}$, (B) $SLN_{1\text{-}26}$ and (C) $SLN_{1\text{-}25}$, respectively.

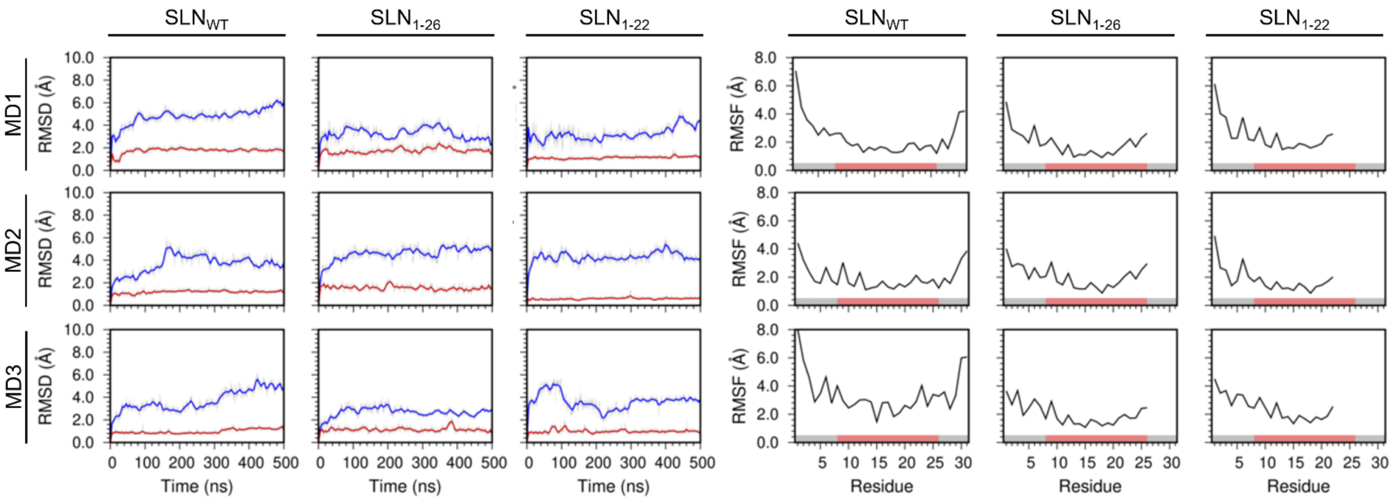


**Figure S19.** Backbone RMSD and per-residue RMSF of $SLN_{WT}$, $SLN_{1\text{-}26}$ and $SLN_{1\text{-}22}$ simulated with the Amber ff14SB force field in the DMPC lipid bilayer. The RMSD of the full-length protein (blue) and the TM domain (red) was calculated by aligning the backbone of the TM domain with the structure at the beginning of each 1-µs MD replicate. RMSF of Cα atoms were calculated from each independent MD trajectory using the backbone of the TM domain as a reference.

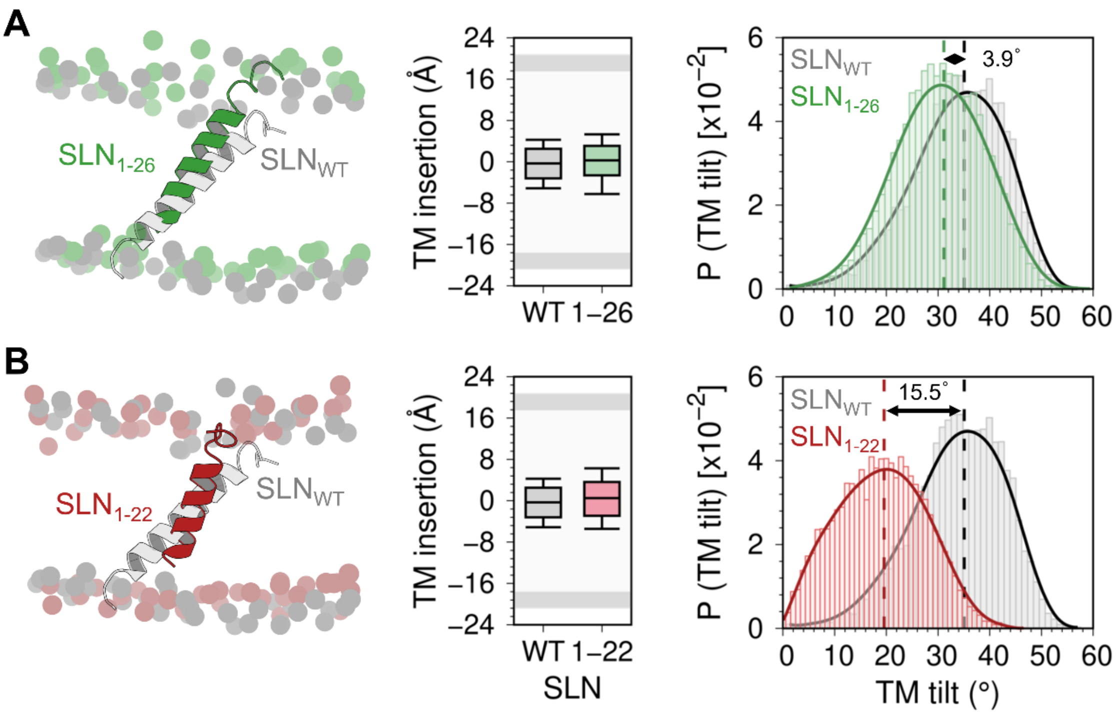


**Figure S20.** Comparison of the structure, TM insertion and tilt angle between $SLN_{WT}$ and (A) $SLN_{1\text{-}26}$ and (B) $SLN_{1\text{-}22}$ simulated with the Amber ff14SB force field in the DMPC lipid bilayer. For comparison, we superposed the most representative structures of $SLN_{WT}$ (gray ribbons) and the SLN deletion constructs (green and yellow ribbons); the lipid heads are show as spheres. Analysis of TM insertion and tilt angle was performed using all six MD trajectories combined; box plots show the full range of variation (from minimum to maximum) using 2.5 %, 50.0 %, and 97.5 % quantiles.

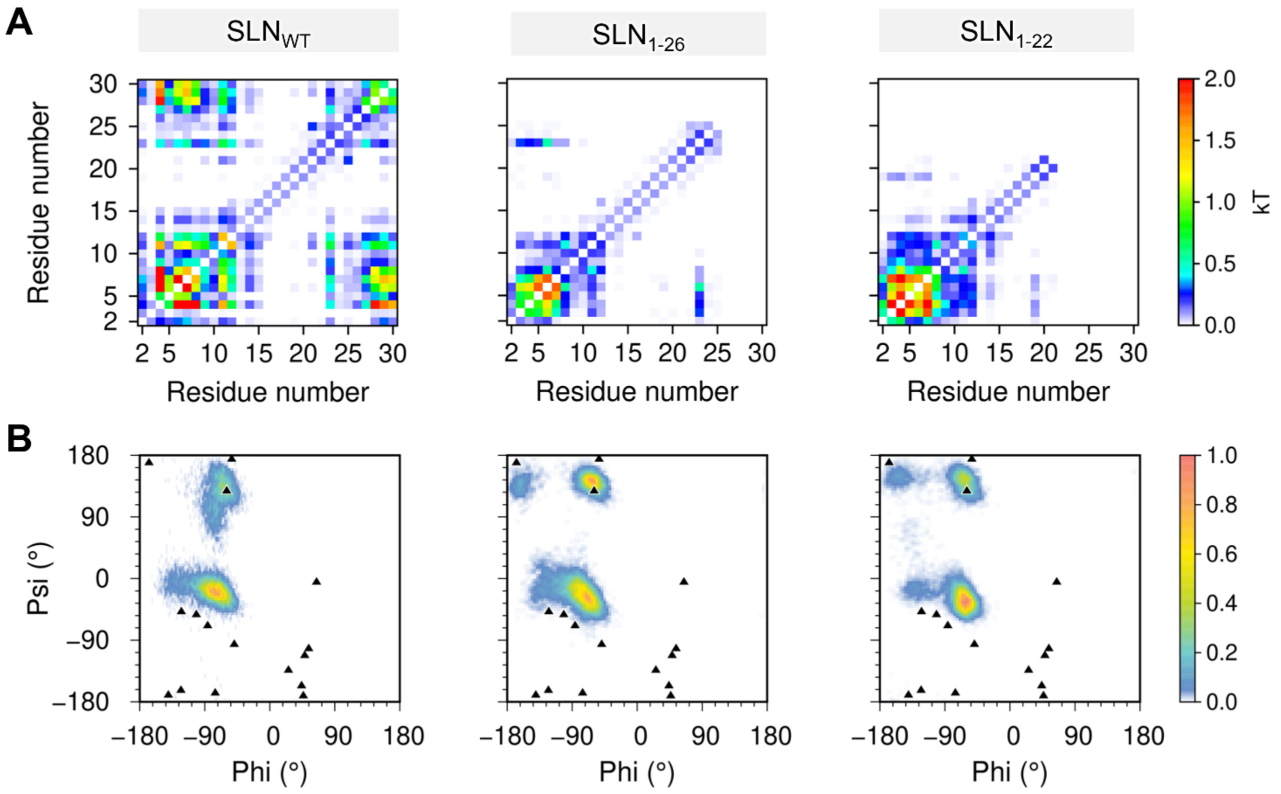


**Figure S21.** Mutual correlation analysis and φ/ψ torsion angle distribution of residue T5 obtained for $SLN_{WT}$ and constructs $SLN_{1-26}$ and $SLN_{1-22}$ simulated with the Amber ff14SB force field in the DMPC lipid bilayer. (A) Mutual information matrix between residues calculated from the trajectories of SLN constructs. (B) Dihedral angle distribution of residue T5, the phosphorylation site of SLN; the φ/ψ torsion angles calculated from the 16 NMR models of SLN (PDB: 1JDM[3]) are shown as black triangles.

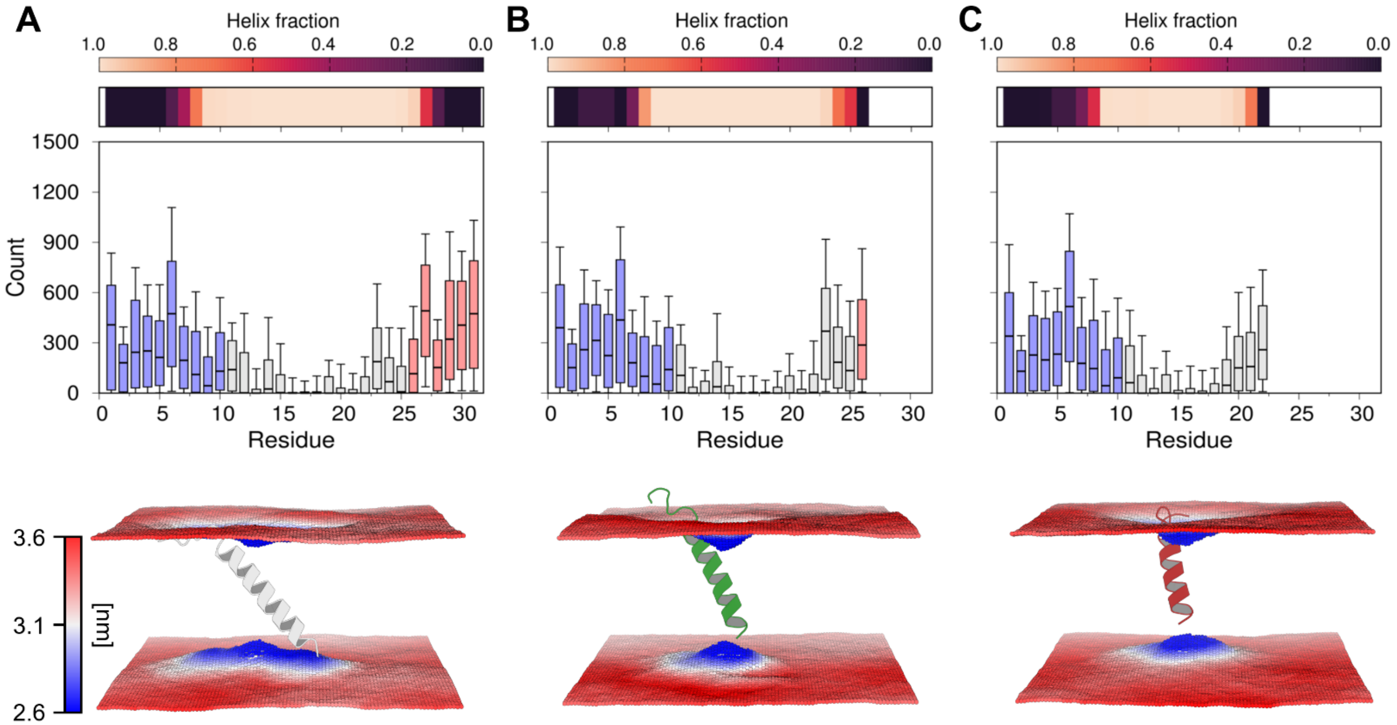


**Figure S22.** Helical content and number of contacts between side chain atoms and lipid headgroup calculated from the SLN constructs simulated with the Amber ff14SB force field in the DMPC lipid bilayer. Local membrane property analysis (bottom) was performed with the *g_lomepro* package.[2] Constructs simulated here are (A) $SLN_{WT}$, (B) $SLN_{1\text{-}26}$ and (C) $SLN_{1\text{-}22}$, respectively.

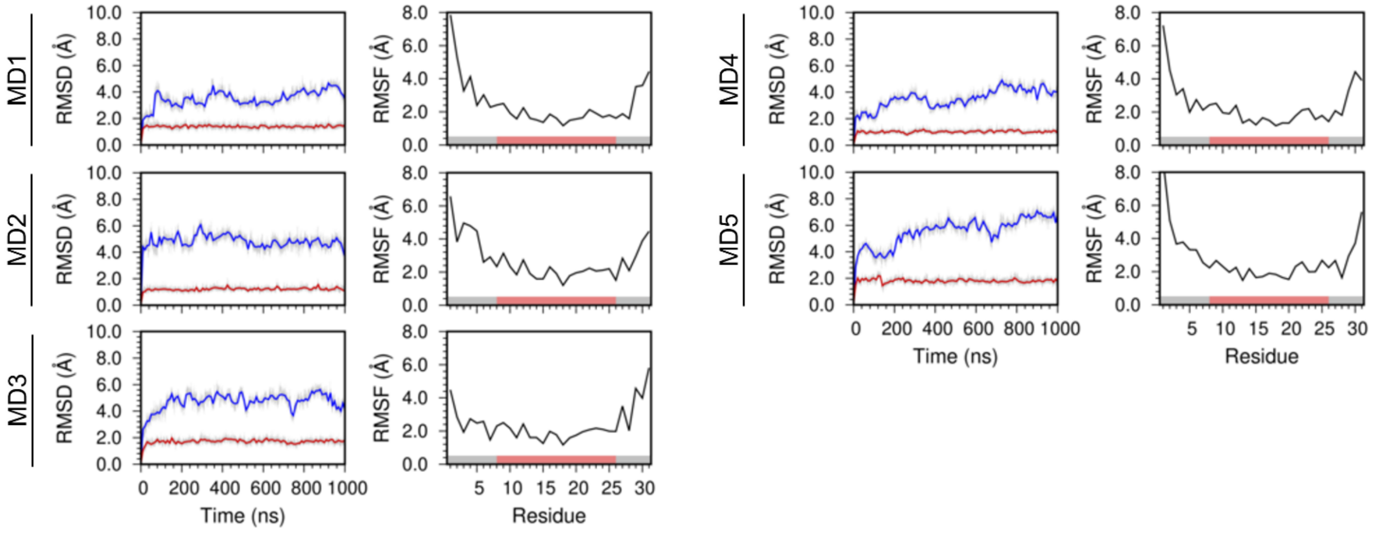


**Figure S23.** Backbone RMSD and per-residue RMSF of SLN phosphorylated at residue T5. The RMSD of the full-length protein (blue) and the TM domain (red) was calculated by aligning the backbone of the TM domain with the structure at the beginning of each 1-μs MD replicate simulated with the Amber ff14SB force field. RMSF of Cα atoms were calculated from each independent MD trajectory using the backbone of the TM domain as a reference.
